# Longitudinal white matter alterations in SIVmac239 infected rhesus monkeys with and without regular cART treatment

**DOI:** 10.1101/2022.05.17.492395

**Authors:** Jiaojiao Liu, Benedictor Alexander Nguchu, Dan Liu, Yu Qi, Alixire, Shuai Han, Yuxun Gao, Xiaoxiao Wang, Hongwei Qiao, Chao Cai, Xiaojie Huang, Hongjun Li

## Abstract

**Objective:** We use the SIV-mac239 infected Chinese rhesus monkeys to longitudinally investigate white matters alterations with and without regular combined antiretroviral therapy (cART) treatment, and its relationship with clinical tests.

**Material and methods:** Diffusion tensor imaging (DTI), CD4 T cell counts, and CD4/CD8 were obtained at baseline, 10 days, 4th week,12th week, 24th week, and 36th week post virus inoculation. postinfection (wpi). Microstructural properties were examined within 76 white matter defined by DTI-WM atlas for rhesus macaques. Corrections for multiple comparisons were performed using a false discovery rate (p < 0.05, FDR). Correlation analyses between imaging markers and clinical measures (CD4 T-cell counts, CD4/CD8 ratio) were determined using Pearson’s correlations.

**Results:** In our model, White matter alterations in SIV-infected macaques can be detected as soon as 4 weeks post inoculation in several brain regions. And with time proceeding, the cART can reverse, relieve, or even progressive effects. CD4 T-cell count is mainly associated with DTI metrics before the cART, whereas CD4/CD8 ratio was associated with white matter alteration with and without cART.

**Conclusion:** SIV-mac239 infection can be an idol modal to explore HIV induced HIV associated brain alterations, and the first group of white matter alterations was as soon as 4 weeks post inoculation in structure next to the periventricular area. As the time progressed, cART can bring different effect to each region, including reversed, relieved, and even progressive effects. In addition, these changes are closely linked to CD4/CD8 ratio even after cART.

**Importance:** 

## 1. Introduction

HIV can shortly invade the central nervous system (CNS) after seroconversion, it can activate microglia and macrophage cells to release toxic factors including excitotoxins, viral products, cytokines and chemokines, resulting in prominent inflammation, immune activation and suppression, and blood–brain barrier disruption, [1–3] which will damage neurons, reflected by cerebral white matter structure alterations. The application of combination antiretroviral therapy (cART), has transformed HIV from a fatal illness into a chronic, manageable condition [4], and may reverse and prevent milder impairments (ANI and MND) to some extent. However, the iatrogenic effects of drugs on brain integrity are also of concern [5,6]. Importantly, previous studies have shown axonal disruption and synaptic injury in HIV patients [7,8]. Diffusion Tensor Imaging (DTI) is a widely used non-invasive neuroimaging technique, which is highly sensitive to microstructural alterations [9]. Previous studies have shown decreased fractional anisotropy (FA), increased mean diffusivity (MD) throughout the brain of HIV-infected patients.[10–21] Even in patients with effective antiretroviral therapies. [11,22] Wherever, there is also study found no significant differences between HIV-infected and healthy controls [23], and some researchers stated that the application of cART did not seem to prevent or reverse existing brain damage. [24–26] However, most the current studies were cross-sectional with several confounders, including but not limited to path of infection, inclusion criteria, duration of infection, complications, different treatment regimens and patient noncompliance, which are difficult to control [13,17,18]. Moreover, the uncertainty of the definite infection data and symptomes, most of the alteration are in months or even years after infection, there is a must for study to investigate early changes in the brain of HIV individual. So longitudinal study with controlled covariations should be conducted to settle the above problem and further to through light upon mechanisms underlying the dynamic alterations with and without the application of cART.

To address the issue, an appropriate animal model with pathological mechanism parallel to HIV patients can be used. Studies have shown that the pathology of simian immunodeficiency virus (SIV)-infected Chinese rhesus monkey was quite similar to that of HIV-infected patients, especially in the disturbance of the central nervous system (CNS), and cognitive or behavioral deficits, and development of AIDS[27–29]. In addition, previous DTI studies on SIV-infected rhesus monkeys have exhibited metric abnormalities similar to that of HIV patients[30,31].

In our present study, we aimed to dynamically examine the white mater alterations of SIV-mac239 infected Chinese rhesus monkeys with and without regularly cART treatment, the relationship between DTI metrics and clinical tests (including CD4 T cell counts, CD4/CD8) of different time points.

## 2. Material and methods

### 2.1 Animal screening prior to main experiments

This study is approved by Beijing Municipal Sciences & Technology Commission. Ten rhesus monkeys were enrolled in the current longitudinal study. Prior to administration to the longitudinal procedure, health screening and indirect immunoinfluscent assay (IFA) were conducted on all the rhesus monkeys to confirm the healthy conditions and exclude the possible infection of simian immunodeficiency (SIV), simian type-D retrovirus (SRV) or simian T-cell lymphotropic virus-I (STLV-I).

### 2.2 Animal model of SIV-infected rhesus monkey and data collection

All the monkeys were inoculated intravenously with SIV-mac239 and were observed regularly. The baseline was defined two weeks before inoculation. Immunological characteristics including peripheral blood CD4^+^, CD8^+^ T-cell countswere collected at the baseline, the 1^st^ week post virus inoculation, the 5^th^ week post virus inoculation, the 12^th^ week post virus inoculation, the 24^th^ week post virus inoculation and the 36^th^ week post virus inoculation. CD4^+^/CD8^+^ ratio were calculated based on the T-cell counts. MRI scans were acquired at the baseline, 10 days post virus inoculation, the 4^th^ week post virus inoculation, the 12^th^ week post virus inoculation, the 24^th^ week post virus inoculation and the 36^th^ week post virus inoculation.

The monkeys were randomly assigned into two groups of therapy group (five monkeys) and control group (five monkeys). All the monkeys were anesthetized by intramuscular injection of ketamine hydrochloride (5-10mg/kg) before each data collection of immunological characteristics, laboratory biochemical characteristics and MRI scan. The five monkeys in therapy group received ART of FTC (50mg/kg/d), TDF (5.1mg/kg/d) and DTG (2.5mg/kg/d) between the third and fourth time points (40 days post virus inoculation).

### 2.3 Housing environment of the subject

All the animals were housed at Institute of laboratory animal sciences, PUMC (Peking Union Medical College) under the same housing standards as that in previous studies [21, 22] including housing temperature maintained at 16~26°C; humidity maintained at 40~70%; 12h/12h light/dark cycle; water provided ad libitum; formula feeds provided on a twice-daily regimen without dietary restrictions.

### 2.3 DTI data analysis

#### 2.3.1 MR protocol

MRI scans were conducted at Beijing YouAn Hospital, Capital Medical University with a 3T Siemens Tim TRIO whole-body magnetic resonance scanner (Siemens, Germany) at each time point to longitudinally assess the impact of SIV infection on brain. The monkeys were anesthetized and placed in the supination position during the MRI scans. T1 weighted images were collected with a turbo flash sequence. The parameters were: repetition time/echo time (TR/TE) =2200/3.54 ms, FOV=128 mm×128 mm, data matrix=192×192, flip angle=9°, slice thickness=1 mm (voxel size=1×1×1.0 mm^2^). DTI images were acquired with a gradient echo single-shot echo planar imaging (EPI). The parameters were: repetition time/echo time (TR/TE) =5200/100ms, FOV=152 mm×152mm, data matrix=76×76, flip angle=90°, slice thickness=2mm (voxel size=2×2×2mm^3^), time points=10min52s.

#### 2.3.2. Macaque Image Processing

Macaque DWI data preprocessing was performed according to previous studies, using the FMRIB Software Library (FSL) (https://fsl.fmrib.ox.ac.uk/) and AFNI (3DSkullStrip for skull-stripping monkey brain data). The DWI raw data preprocessing was conducted to correct eddy currents, susceptibility-induced distortions, and animal movements. The data were then fitted using DIT-tensor fitting technique available in FSL. Four types of diffusivity maps were generated: fractional anisotropy (FA), a measure of the directionality of water diffusion; mean diffusivity (MD), a measure of the water diffusivity in the transverse directions[32](Chang et al., 2020). Each of these macaque’s diffusivity maps(FA, MD) was registered to the standard space template, the diffusion-tensor-based white matter atlas for rhesus macaques (also known as UWDTIRhesusWMAtlas, https://www.nitrc.org/projects/rmdtitemplate/)[33], using FMRIB’s Linear/Non-linear Image Registration Tools (FLIRT/FNIRT), part of the FSL version 5.09 [34]. The template is population-based developed from a large number of animal high-quality scans (N=271) that allows it to account for variability in the species; It has a high signal-to-noise ratio (SNR) and FA values, and high image sharpness with visible small white matter structures and spatial features [33]. Two neurologists (X.W. and B.N., with 8 and 4 years of experience, respectively) inspected the data visually to confirm the registration accuracy

#### 2.3.3. Regions of Interest (ROIs) Selection and WM Microstructural Property Analysis

Microstructural properties were examined within 76 white matter defined by DTI-WM atlas for rhesus macaques. All WM structures were assessed in our analyses. Each WM region of interest (ROI) was analyzed longitudinally for changes across time. These microstructural changes were examined at baseline and after at baseline, 10 days, 4 weeks, 12 weeks, 24 weeks, and 36 weeks post virus inoculation.

#### 2.3.4. Statistical analyses

Statistical analyses were performed on SPSS software [IBM SPSS Statistics for Windows, version 20.0 (IBM Corporation, Armonk, New York, USA)]. Group differences between Macaque having cART and those without cART, the effect of time on microstructural properties, and the interaction effect of cART treatment and time were determined using the Two-factor mixed-design ANOVA. Of 10 monkeys infected with HIV, 5 were randomly assigned to undergo cART treatment and the other 5 received no treatment. Hence, the between-subjects factor had two levels. Over 36 week period, microstructural properties were measured at six time points, representing six levels of the “within-subjects” factor. These levels were: the baseline [time point 1], after 10 days [time point 2], 4 weeks [time point 3], 12 weeks [time point 4], 24 weeks [time point-5], and 36 weeks, at the end of the programme [time point-6].

Corrections for multiple comparisons were performed using a false discovery rate (p < 0.05, FDR). Correlation analyses between imaging markers and clinical measures (CSF Viral Load, Plasma Viral Load, CD4 T-cell counts, CD8 T-cell counts, CD4/CD8 ratio) were determined using Pearson’s correlations.

## Results

### *Longitudinal DTI alterations in macaque without treatment* [Figure 2]

**Figure 1.**
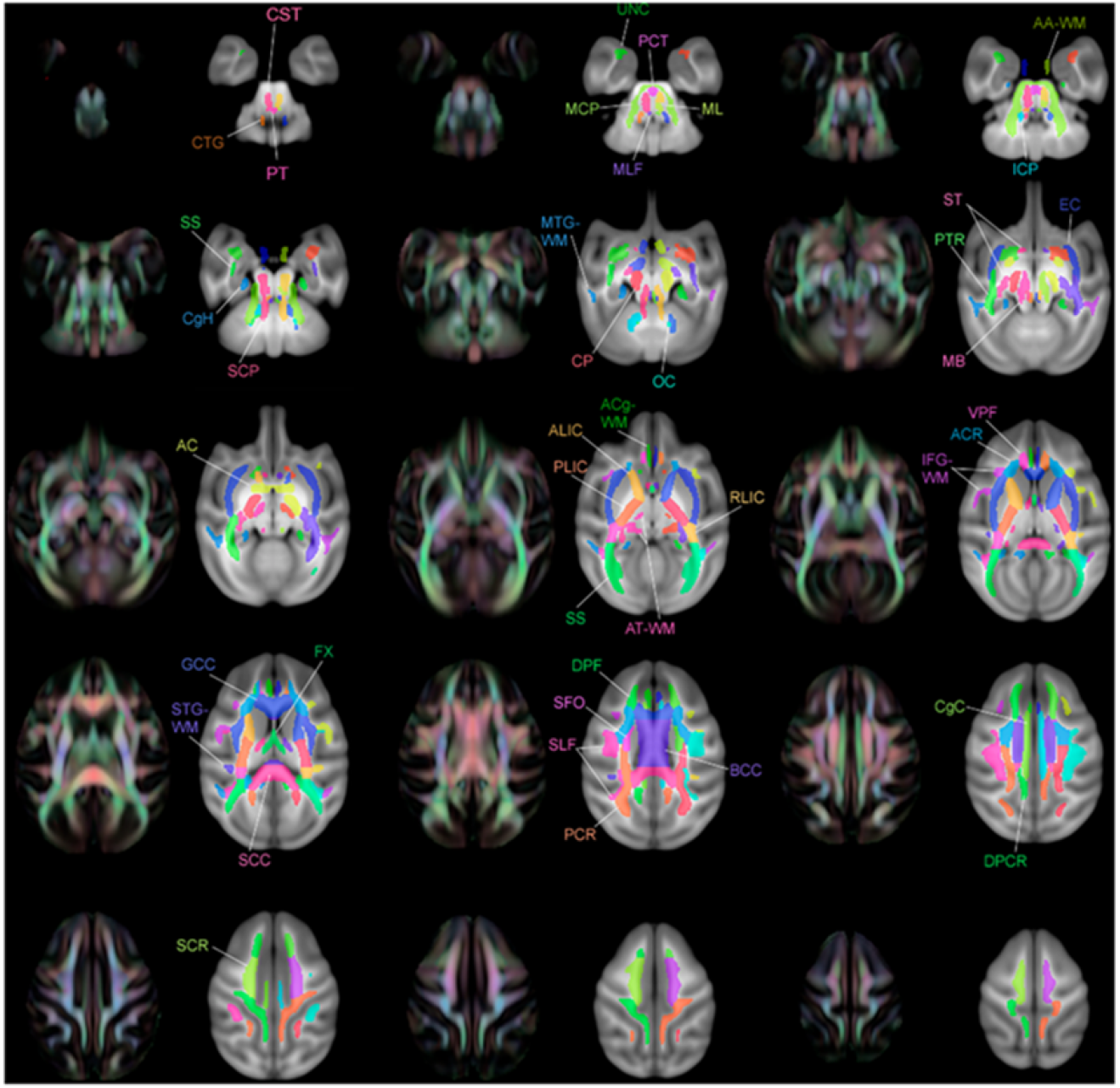
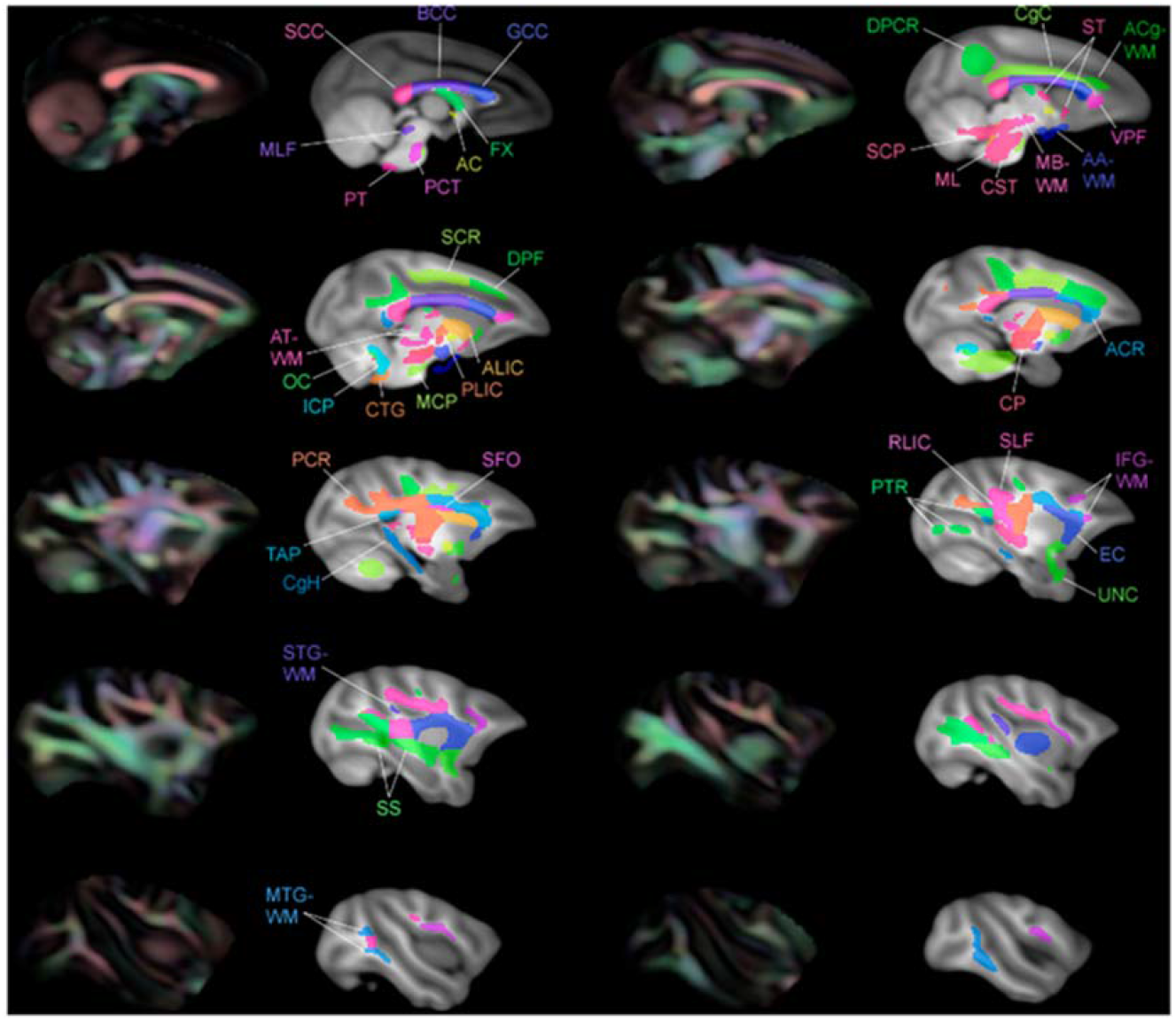
Multiple slices of atlas overlaid on T1-W template illustrating the regions of interest (ROIs) beside FA color image. These ROI images were adapted from Zakszewski et al. [33]

**Figure 2.**
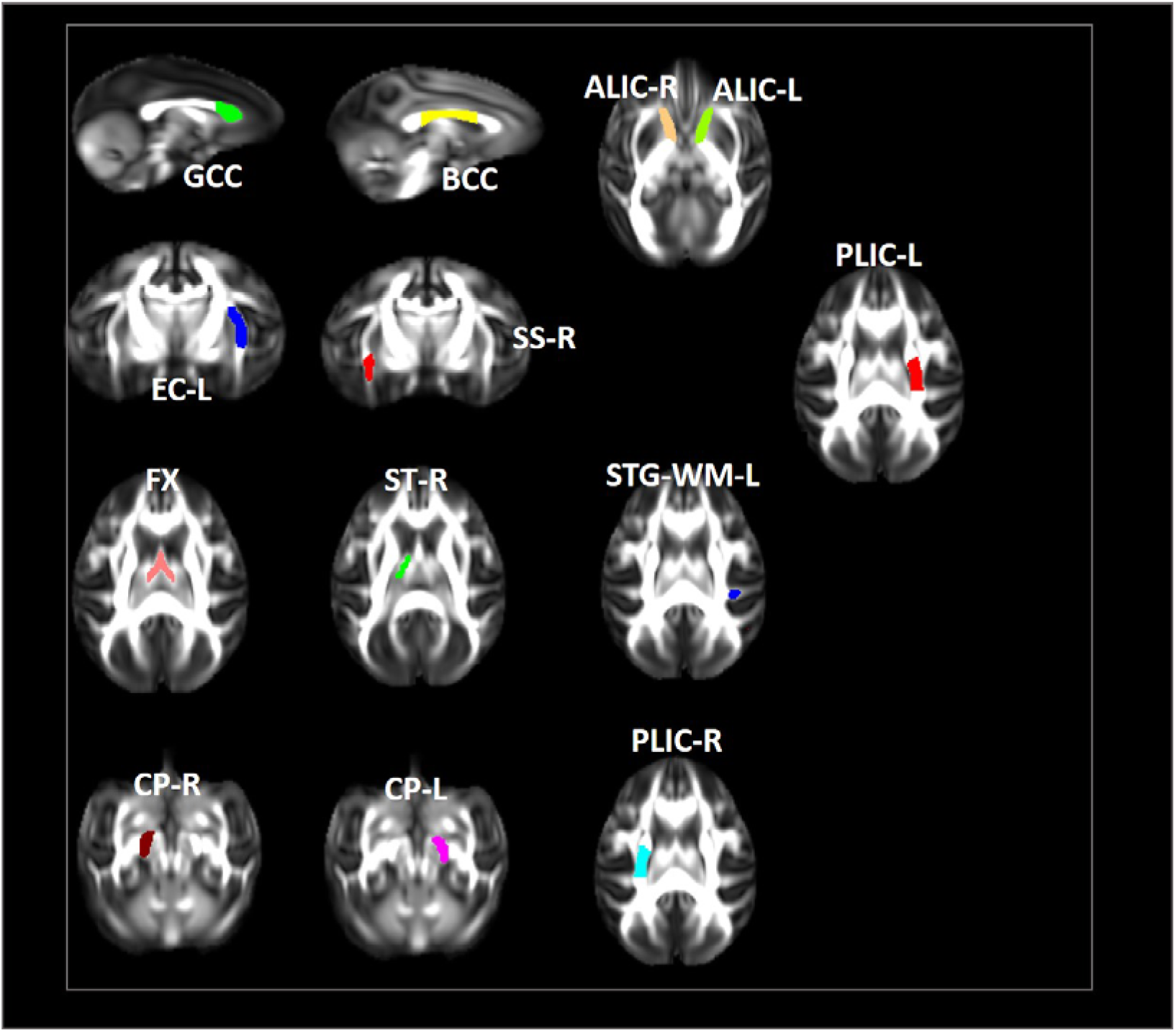
Changes in microstructures propagate differentially in SIV-macaques with and without antiretroviral therapy.

**Fig3.**
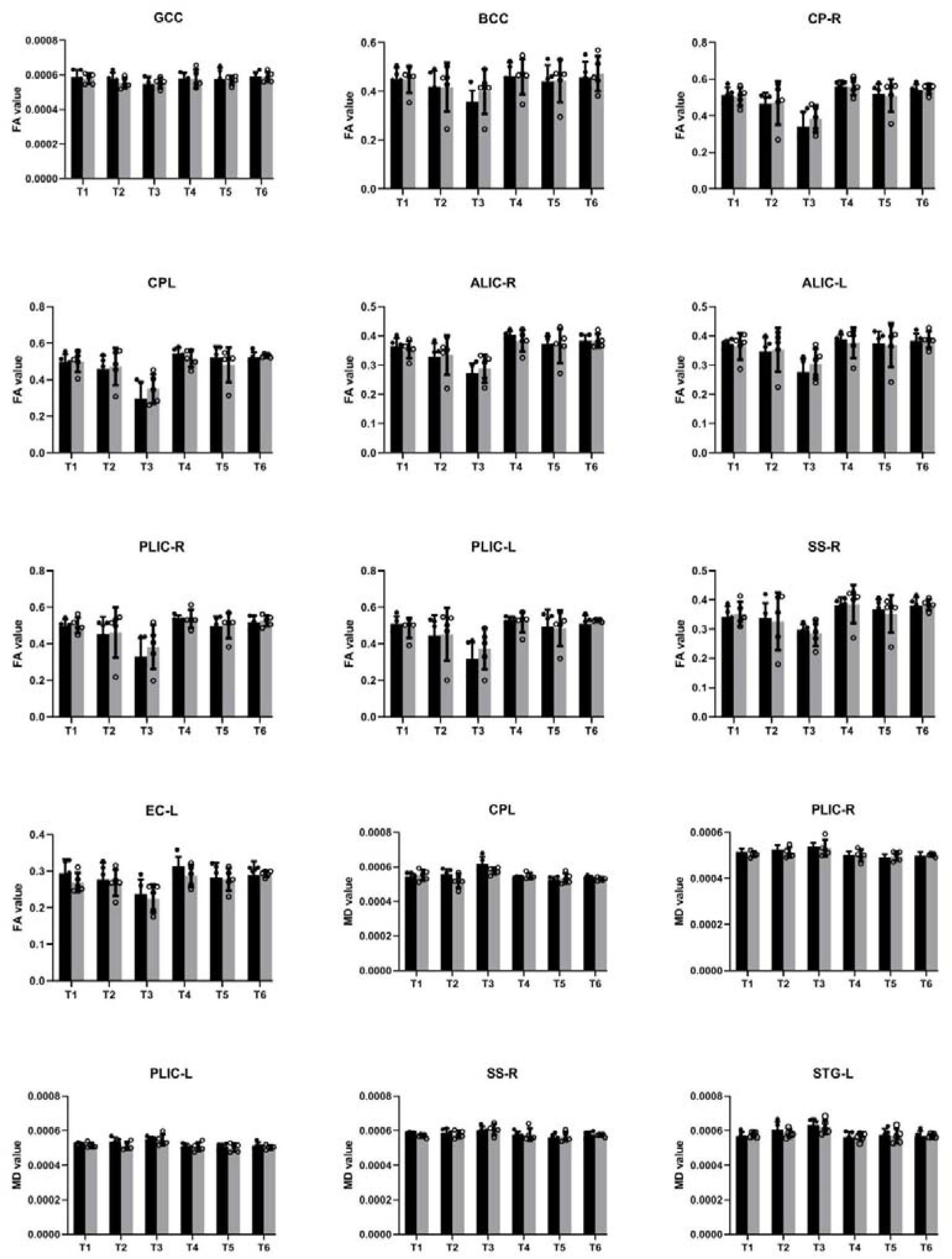
Illustration of significant alterations of FA and MD across time incART+ andcART+ groups. Bars, mean ± SE. Black, SIV-mac239 infected macaques without treatment; Gray, SIV-mac239 infected macaques with regular treatment. Note: T1 the baseline; T2, 10 days post virus inoculation; T3, the 4th week post virus inoculation; T4, the 12th week post virus inoculation; T5, the 24th week post virus inoculation; T6, the 36th week post virus inoculation.

In our study, we longitudinally study the white matter alteration from the baseline, 10 days post inoculation, 4 weeks post inoculation, 12 weeks post inoculation the infection, 24 weeks post inoculation the infection, and 36 weeks post inoculation. We found that the alteration was firstly examined in 4 weeks after inoculation insuperior temporal gyrus (STG) (*p*=0.029244), posterior limb of internal capsule (left) (PLIC-L) (p=0.023222), fornix(FX) (*p*=0.041225), cerebral peduncle left (*p*=0.0173969), cerebral peduncle right (*p*=0.024109), left anterior limb of internal capsule(ALIC-L) (p= 0.0162715), right anterior limb of internal capsule(ALIC-R) (p= 0.01556)and right sagittal striatum(SS-R) (p= 0.0190229) on FA, and adjacent amygdala white matter (AA-WM-RR)(*p*=0.004665), posterior limb of internal capsule -left (PLIC-L) (p= 0.0232217) (*p*= 0.0451514)and right sagittal striatum (SS-R) (*p*= 0.0292208)on MD. While 24 weeks post inoculation we found the alterations were then appeared on the regions of posterior limb of the internal capsule-right(PLIC-R) (*p*= 0.0426243), genu of corpus callosum (GCC) (*p*=0. 0.0192415),body of corpus callosum (BCC) (*p*=0.032449), external capsule-left (EC-L) (*p*= 0.01540547) (*p*= 0.0447482), striaterminalus-right (ST-R) on FA(*p*= 0.0029465), and PLIC-R(*p*= 0.016532) (*p*= 0.0328549) on MD.

### Longitudinal DTI alterations in macaque on regular cART

After cART, we found that there were no alterations in BCC, FX for FA and MD, no progression in AA-WM-R for MD, and PLIC-R, PLIC-L for FA, progression was slow in STG-WM-L, GCC, CP-R, CP-L, no effect in PLICR, ALIC-R, SS-R, ST-R. Alternatively, in EC-L.

### Correlation between Altered DTI properties on regular cART and clinical metrics[Table 1]

**Table1.** The correlations between DTI metrics and clinical tests with regularcART treatment.

| Target region |  | CD4 |  | CD4/CD8 |  |
| --- | --- | --- | --- | --- | --- |
|  |  | r | p | r | p |
| MD | GCC | NA | NA | -0.8852612 | 0.045843 |
|  | ALIC-R | NA | NA | -0.86564256 | 0.057912 |
|  | PLIC-L | NA | NA | -0.94726 | 0.0144217 |
|  | ALIC-L | NA | NA | -0.891353 | 0.0422815 |
|  | ST-R | NA | NA | 0.859125 | 0.06211 |
|  | AA-WM-R | NA | NA | 0.90225 | 0.026141 |
| FA | CP-R | NA | NA | 0.9260468 | 0.0238721 |
|  | CP-L | NA | NA | 0.900711 | 0.0369919 |

Note: NA means without significant correlation. GCC, Genu of Corpus Callosum; ALIC-R, Anterior Limb of the Internal Capsule - Right ; PLIC-L, Posterior Limb of the Internal Capsule - Left ; ALIC-L, Anterior Limb of the Internal Capsule - Left; ST-R, Stria Terminalis - Right ; AA-WM-R, Adjacent Amygdala White Matter - Right; CP-R, Cerebral Peduncle - Right; CP-L, Cerebral Peduncle - Left..

We examined the relationship between clinical measures, such as CD4+ T cell counts and CD4/CD8 ratio, and the alterated DTI metrics with great significance in cART-treated macaques. We confirmed that increased MD and FA are with decreased CD4+T cell counts in Only one region in AA-WM-R (p=0.073927, R=-0.84149). And positive relation only for ST-R (p= 0.06211, R= 0.859125). Also, for CD4/CD8 ratio, we found increased MD is with low CD4/CD8 ratio in GCC (p=0.045843, R=-0.8852612), ALIC-R (p= 0.057912, R= −0.86564256), PLIC-L (p= 0.0144217, R=-0.94726), ALIC-L (p= 0.0422815, R= −0.891353), and positive relations in ST-R(p= 0.06211, R= 0.859125) and AA-WM-R(p=0.026141, R=0.90225), with regularcART treatment. As for FA, CP-R(p= 0.0238721, R= 0.9260468) and CP-L(p= 0.0369919, R= 0.900711) (p= 0.0309355, R= 0.91197044) showed positive relation.

### Correlation between Altered DTI properties without treatment and clinical metrics[Table 2]

**Table2.** The correlations between DTI metrics and clinical tests without treatment.

| Target region |  | CD4 |  | CD4/CD8 |  |
| --- | --- | --- | --- | --- | --- |
|  |  | r | p | r | p |
| MD | PLIC-R | -0.933691 | 0.0202919 | NA | NA |
|  | PLIC-L | -0.86411685 | 0.0588879 | NA | NA |
|  | STG-WM-L | 0.9826859 | 0.0027277 | NA | NA |
|  | PLIC-R | NA | NA | -0.93372 | 0.02027519 |
|  | FX | NA | NA | -0.8782170 | 0.0500745 |
|  | ST-R | NA | NA | -0.865228 | 0.058177 |
| FA | PLIC-L | 0.946059 | 0.014916 | NA | NA |
|  | PLIC-R | 0.9057661 | 0.034230 | NA | NA |
|  | CP-R | 0.94168096 | 0.0167576 | NA | NA |

Note: NA means without significant correlation. PLIC-R, Posterior Limb of the Internal Capsule –
Right; PLIC-L, Posterior Limb of the Internal Capsule – Left; STG-WM-L, Superior Temporal Gyrus WM – Left ; PLIC-R, Posterior Limb of the Internal Capsule – Right; FX, Fornix; ST-R, Stria Terminalis – Right; CP-R, Cerebral Peduncle – Right.

We also estimated the correlation between clinical measures, such as CD4+ T cell counts and CD4/CD8 ratio, and the alterated DTI metrics with great significance. We found that decreased FA was with decreased CD4+ T cell counts in PLIC-L (*p*=0.014916, R=0.946059), PLIC-R (t1, *p*=0.034230, R=0.9057661), CP-R (p= 0.0167576, R= 0.94168096) in monkeys without treatment. Increased MD is with low CD4+ T cell counts in PLIC-R(p=0.0202919, R= −0.933691) (*p*= 0.0517050, R= −0.87555), and PLIC-L (p=0.0588879, R=-0.86411685), except for the positive relation STG-WM-L(p= 0.0027277, R= 0.9826859). As for CD4/CD8 ratio, increased MD was with decreased CD4/CD8 in PLIC-R(p= 0.02027519, R= −0.93372), FX (t1, p= 0.0500745, R= −0.8782170), and ST-R(p= 0.058177, R= −0.865228).

## Discussion

In the current study, longitudinal cerebral white matter structure alterations were assessed in 10 SIV-mac239 infected monkeys with and without regular cART treatment. We found significant white matter structure alterations in these monkeys, consisted of higher mean diffusion and lower fractional anisotropy, as conducted by DTI. TBSS showed that these effects were progressive with time. In addition, we found that the disturbance of white matter was appeared as early as 4 weeks post inoculation, and that with regular cART the damage can be alleviated or even reversed. Furthermore, DTI metric alterations were significantly correlated with clinical measures such as CD4 T cell counts, CD4/CD8.

The underlying etiology of white matter alterations in HIV individuals is still unclear. Possible pathophysiologic mechanism include but not limit to synaptodendritic injury, inflammation, demyelination, and even microvascular abonormalitis correlated to concomitant cardiovascular risk factors, especially hypertension. In the context of HIV based on DTI acquisition parameters, FA reflects the aspect of axonal structural integrity reflecting cytoskeleton and membrane integrity[35] and proposed structural integrity ofrocytic[36]. MD was interpreted as related to the degree of WM microstructure density[35, 37], such as extracellular space between WM tracts[36] that may be affected by neuroinflammation including glial involvement and myelin loss to a lesser extent. In this case, myelin contribution to MD changes is debatable[36] as massive demyelination is not a common feature of HIV-related brain injury even in those who had severe HIV encephalopathy[38] Previous studies on HIV patients and matched healthy controls have shown subtle but widespread white matter alterations, especially findings of alterations in fractional anisotropy and mean diffusion [39–49], suggesting disorganization of the micro- and macro-structure of the white matter. [50]

However, few studies have explored the original time and first groups of regions that the white matter have been destroyed, and what happened after early cART (as early as 40 days post inoculation), all of which can throw light upon the underlying neuropathological mechanism, and illustrate the effect of timely cART on imaging findings, immune indicators and the relationship between them. In our study, we longitudinally study the white matter alteration of rhesus monkeys with and without regular cART treatment from the baseline, 10 days, 4 weeks, 12 weeks, 24 weeks, and 36 weeks post inoculation. In our present study, we found that the alteration was firstly examined in 4 weeks after inoculation in Amygdala, superior temporal gyrus (STG), posterior limb of internal capsule (left), fornix, bilateral cerebral peduncle, and bilateral anterior limb of internal capsule on FA, and posterior limb of internal capsule -left (PLIC-L) and right sagittal striatum (SS-R) on MD, which suggest that these regions are more sensitive to SIV infection. Studies have shown that HIV enters the central nervous system via monocytes and perivascular macrophages very early after seroconversion. The virus then replicates and induces neuronal damage mainly by neuroinflammatory mechanisms triggered by infected microglial cells, and the inflammation selectively occurs in the dopamine-rich areas of the brain including the subcortical areas, particularly the dopamine-rich basal ganglia and induces a subcortical dementia. [51–54] Our findings of alternated FA in Amygdala and sagittal striatum, which are parts of the basal ganglia, is consistent with this body of work. The fornix is a major efferent tract of the hippocampus, a structure critical for normal memory function, which has been reported on AD, schizophrenia, and multiple sclerosis. Microstructural alteration of the fornix is a contributor to early episodic memory dysfunction in non-demented individuals[55–58]. Internal capsule is a region that is sensitive to WM damage in those with cognitive impairment and AIDS. [59] [78]Previous study also reported that the higher HIV RNA density of infected macrophages was mostly in the subcorticle white matter including IC. [60] In addition, many studies have also reported white matter injury (i.e., decreased FA or increased MD) in the subcortical white matter, particularly of the internal capsule. [61–65] As for the alteration of cerebellar peduncle and sagittal stratum, previous Study on adolescent had obtained similar changes[66].

While in 12 weeks post inoculation, regions of PLIC-R, GCC, BCC, EC-L, and ST-R showed significant changes on FA, and PLIC-R on MD. Early cerebral infiltration of HIV can occur before antiretroviral medications are administered, exposing the surrounding white matter to the neurotoxic effects before effective viral suppression. It has been suggested that the periventricular white matter such as CC is especially vulnerable to viral attack owing to its proximity to cerebrospinal fluid, which is an HIV reservoir that can carry the virus within 8 days of infection[67,68,69]. Studies also showed that the highest density of HIV infected macrophages was in the subcorticle white matter including CC. [70] Consistent with this early impact on the brain, white matter degradation has been found within the first 100 days after HIV transmission[71]. As a major communication pathway between the hemispheres, the CC is responsible for the functional integration of complex cognitive, motor, and behavioral tasks. So the observation of CC can be an indicator of underlying neurocognitive disorder of HIV individuals. External capsule serves as pathway for psychomotor functions, many previous studies have shown alternated external capsule in HIV patients. [72,73]

After cART, we found that initial alteration in BCC and FX disappeared, no progression in AA-WM-R, PLIC-R, PLIC-L, progression was slow in STG-WM-L, GCC, CP-R, CP-L, no effect in PLICR, ALIC-R, SS-R, ST-R. Alternatively, in EC-L, we found that cART can damage the structure to some extent. White mattet repair afforded by viral suppression and cART could lead to greater degree of axonal structural integrity that is above normal effects of aging. This may be particularly possible in the internal capsule as it is a region that is sensitive to white matter damage in those with cognitive impairment and AIDS. [59] The lack of aggressive HIV associated WM damage in the CC after cART suggested that early treatment is neuroprotective to the CC to some extent. [74] On one hand, cART may contribute to synaptic injury via oxidative stress as has been demonstrated in vitro and in animal models[75]. On the other hand, early and continuous antiretroviral therapy can be neuroprotective, limiting the damage to white matter. [76,77]

In our study, we found that decreased CD4+ T cell counts were with decreased FA value and increased MD value in patients without cART treatment, however after early and regular cART treatment, there was no significant changes between CD4 T cell counts and DTI metrics, indicating that absolute CD4 T cell counts may fail to accurately reflect the risks threating HIV individuals since immune dysfunction persists with normalization of CD4 counts. [78] In addition, we found that increased MD was with decreased CD4/CD8 in many brain regions, including GCC, ALIC-R, PLIC-L, ALIC-L, PLIC-R, and FX, in macaques with and without regular cART treatment. To our best knowledge, the lower CD4+/CD8+ ratio could be attributed to the persistent inflammation and immunosenescence caused by viral infection[79] and can also be used as a biomarker for T-cell activation to characterize the migration of T cells into the CNS after HIV infection and the production of inflammatory cytokines, which can indirectly lead to white matter damage. Our result indicated that in the cART eara, CD4/CD8 maybe a more sensitive clinical biomarker for assessing risks facing the modern aviremic HIV population. [80]

## Limitations and further consideration

Firstly, our study failed to record cognitive performance, for it takes a lot of time to train the monkeys for specified tasks, which limited us to explore the underlying mechanism of neurocognitive dysfuctions and the correlations among them with DTI metrics and clinical tests. In addition, the small sample size limited us for effective statistic analyses. So in the future study, we would include a lager sample of macques.

## In conclusion

SIV-mac239 infection can be an idol modal to explore HIV induced HIV associated brain alterations, and the first group of white matter alterations was as soon as 4 weeks post inoculation in structure next to the periventricular area. And alterations progressed with time proceeding, cART can bring different effects to each region, including reversed, relieved, and even progressive effects. In addition, these changes are closely linked to CD4/CD8 ratio even after cART. Further studies are in great need to illustrate underlying mechanism behind them.

## Ethic statement

The study was approved by the Institutional Animal Care and Use Committee (IACUC) at the Institute of Laboratory Animal Science, Chinese Academy of Medical Sciences (IACUC Approval No: LHJ18001), and performed according to the recommendations in the Guide for the Care and Use of Laboratory Animals of the Institute of Laboratory Animal Science and the recommendations of the Weatherall report for the use of nonhuman primates in research (http://www.acmedsci.ac.uk/more/news/the-use-of-nonhuman-primatesin-research/) to ensure personal safety and animal welfare. All macaques were housed and fed in an Association for Assessment and Accreditation of Laboratory Animal Care (AAALAC)-accredited bio-safety level 3 laboratory.

## Data and materials availability

All data are available in the main text.

## Acknowledgements

This work was supported by the National Natural Science Foundation of China (grant nos. 61936013); the Beijing Natural Science Foundation (7212051).

